# Spatiotemporal variation in abundance and genetic structure across the urban-rural landscape gradient: *Aedes albopictus* (Skuse, 1894) (Diptera: Culicidae) in Wake County, NC

**DOI:** 10.1101/2025.06.12.659333

**Authors:** Emily MX Reed, Michael H Reiskind, Martha O Burford Reiskind

## Abstract

Since its invasion of the United States in the 1980s, *Aedes albopictus* (Skuse, 1894) has become a major pest and a significant public health threat in the Southeastern USA. Despite its importance, we know little about its population genetics at fine spatial scales. To remedy this lack of information, we analyzed *Ae. albopictus* spatial variation in mosquito abundance and genetic structure in an urban-rural landscape over two years (2016 and 2018) in Wake County, NC, USA. We used a reduced representation sequencing method to generate between 11,00 to 30,000 SNPs that were used for population genetic analyses. We found spatial variation in both the abundance, genetic diversity, and significant differences in genetic divergence among sites that varied between the two years. The year-to-year variation in the population genetic patterns at the within county scale suggests a dynamic system which requires extensive geographic, temporal, and genomic sampling to resolve.

**Graphical Abstract:** 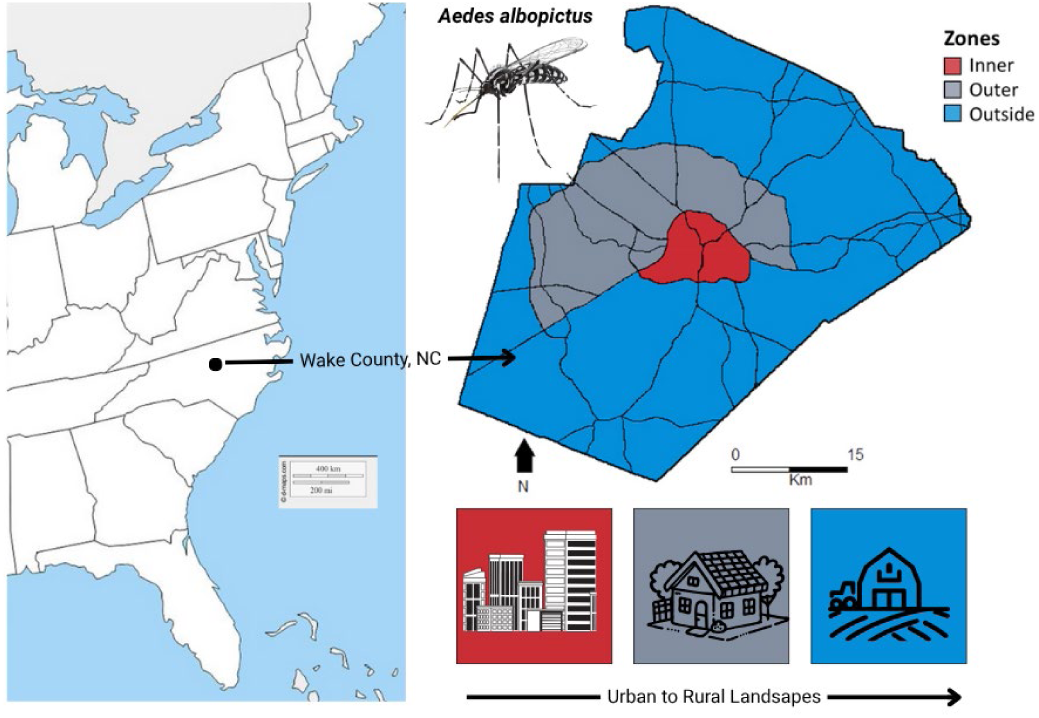

## Introduction

*Aedes albopictus* is a cosmopolitan mosquito that is invasive in most of its range outside of east Asia, spanning a large latitudinal range under current climate conditions, from the Equator to 50°. This mosquito prefers suburban environments, where it threatens human and wildlife health as a vector of viral and filarial pathogens (Hawley et al. 1987, Gratz 2004, Barker et al. 2003, Spence Beaulieu et al. 2019). This is of concern as changes in global temperatures and land use may allow *Ae. albopictus* to both invade new temperate regions, especially in warmer urban landscapes, and increase the risk of human and animal disease (Gratz et al 2004). Given its historic arrival in the Southeastern USA (in 1985, Hawley et al. 1987) and more recent expansion into the northeastern USA (Hahn et al. 2017) and California (Metzger et al. 2017), we need a clear understanding of the behavior and movement of the species along urban rural gradients and within urban settings.

Landscape, the spatial pattern of suitable habitat patches and unsuitable matrix, and human population density help determine the distribution, abundance, age structure, and genetic diversity of species. These demographic characteristics can be altered by land use change, such as urbanization. Urban landscapes are mosaics of habitat patches, such as parks, forests, and the built environment, and are characterized by anthropogenic impacts on abiotic conditions such as temperature, habitat fragmentation, and increased impervious surface cover (Vitousek 1997, Irwin and Bockstael 2007, Baudouin et al. 2018). In conjunction with habitat alternations, the pattern in human population density and energy consumption or “urbanization” is referred to as the urban-rural gradient (McDonnell & Pickett 1990). From an ecological perspective, urbanization and the urban-rural gradient describes the gradual reduction in human population density and intensity of its influence on the environment (Herrero-Jáuregui et al. 2018). As such, this gradient predicts that population abundance of native species is inversely related to degree of urbanization, and that population genetic differentiation increases with urbanization (e.g. McKinney 2002, Johnson and Munshi-South 2017). In native species this can cause a loss of genetic diversity and higher differentiation in urban populations (e.g. native cricket in urban environments; Vandergast et al. 2008). However, species response to urbanization is variable and depends both on the ecology of the taxa, the urban area(s), and the surrounding landscape (McIntyre 2000, McDonnell and Hahs 2008). For human dependent, invasive species there may be a pattern of greater abundance in urban centers and potential for greater genetic diversity (Reed et al. 2020). The patterns of abundance and population genetic structure of public health threats like *Ae. albopictus* are critical to document along urban—rural landscape networks (McDonnell et al. 1997, Blair and Johnson 2008, Ariori et al. 2017).

Much of *Ae. albopictus’s* invasion success can be attributed to its ecology. *Aedes* mosquitoes in the subgenus *Stegomyia* lay desiccant-resistant eggs in small, ephemeral pools of water. Containers used for oviposition can be natural (e.g. tree holes, bamboo shoots) or anthropophilic (e.g. planters, clogged gutters, bird baths, and human refuse). These artificial containers are commonly exploited by *A. albopictus* and have contributed to its spread along with international trade (Hawley et al. 1987). *Aedes albopictus* larvae may also have a competitive advantage over other mosquito species, contributing to its invasion success (Juliano 2010). The global distribution of *Ae. albopictus* is limited primarily by climate (Benedict et al. 2007, Cunze et al. 2016, Ding et al. 2018, LaPorta et al. 2023). At finer spatial scales, *Ae. albopictus* presence, abundance, and growth depends on microclimate and landscape features, including urbanization (Medley et al 2021, Murdock et al. 2017, Spence Beaulieu et al. 2019, Hopperstad et al. 2020, Reiskind et al. 2017). *Aedes albopictus* abundances can also vary widely over relatively short time periods, making it difficult to estimate true population sizes in an area and disentangle the relative importance of spatial and temporal variation (Crocker et al. 2017). Population genetic studies of *Ae. albopictus* in eastern North America have suggested significant population structure at coarse scales (Stone et al. 2020, Gloria-Soria et al 2023).

To better understand how local *Ae. albopictus* abundances, stability, and connectedness vary across landscapes, we used both abundance and genetic data to describe population-level patterns in Wake County, North Carolina, USA. *Aedes albopictus* was first recorded in Wake County in 1993 (Kraemer et al. 2015, Kraemer et al. 2017). Its congener *Ae. aegypti* has not been detected in North Carolina since 1995, shortly after *Ae. albopictus*’ arrival (Reed et al. 2019, Reiskind et al. 2020). *Aedes albopictus* in Wake County are most active during the summer and enter diapause during the winter (Reed et al. 2019). Surveys of *Ae. albopictus* and other container breeding mosquitoes have been conducted in Wake County (Reed et al. 2019, Spence-Beaulieu et al. 2019, Reiskind et al. 2020). These studies primarily focused on climate variables, and when land-use variables were investigated, researchers found only marginal correlations with *Ae. albopictus* abundances. However, several factors may have influenced these studies, including sampling methods and the area of land around a collection site used to quantify landscape predictors.

In this study, we analyzed the population abundance and genetic characteristics of *Ae. albopictus* in Wake County. We used abundance and genomic data from preserved *Ae. albopictus* individuals from a 2016 Wake County survey (Reed et al. 2019), reared from eggs collected in ovitraps, and from adult traps from more locations in 2018. We looked broadly at patterns within and between designated “zones” of urbanization intensity. Integrating genomic methods with surveillance further elucidates the effects of landscape and urbanization on the population dynamics of invasive species at fine spatial scales and provides place-based information on *Ae. albopictus* population structure critical for vector control.

## Methods

### Study Site

Wake County area is 2,220 Km^2^, has a population of 1.1 million in 2019, and an average population density of 495.5 people/ Km^2^ at the time of this study. Urban areas of Wake County are patchily distributed around Raleigh, its largest and most central city. To compare differences along the urban—rural gradient, we grouped sites in each county into geographically-delineated “zones,” corresponding to differing levels of urbanization and settlement patterns (Fig. 1). Urbanization is highest in the center of Wake County, where the city of Raleigh is located, and transitions into suburban and rural areas on the outskirts of the county. Therefore, we grouped sites from Wake County into three zones separated by the major highways surrounding Raleigh: (1) Inner zone within the interstate I-440 and Interstate 40 beltway (urban to suburban), (2) Outer zone between I-440, the I-540 beltway (suburban to rural), and US route 1, and (3) Outside zone as the remainder of Wake County (suburban to rural) (Fig. 1).

**Figure 1.**
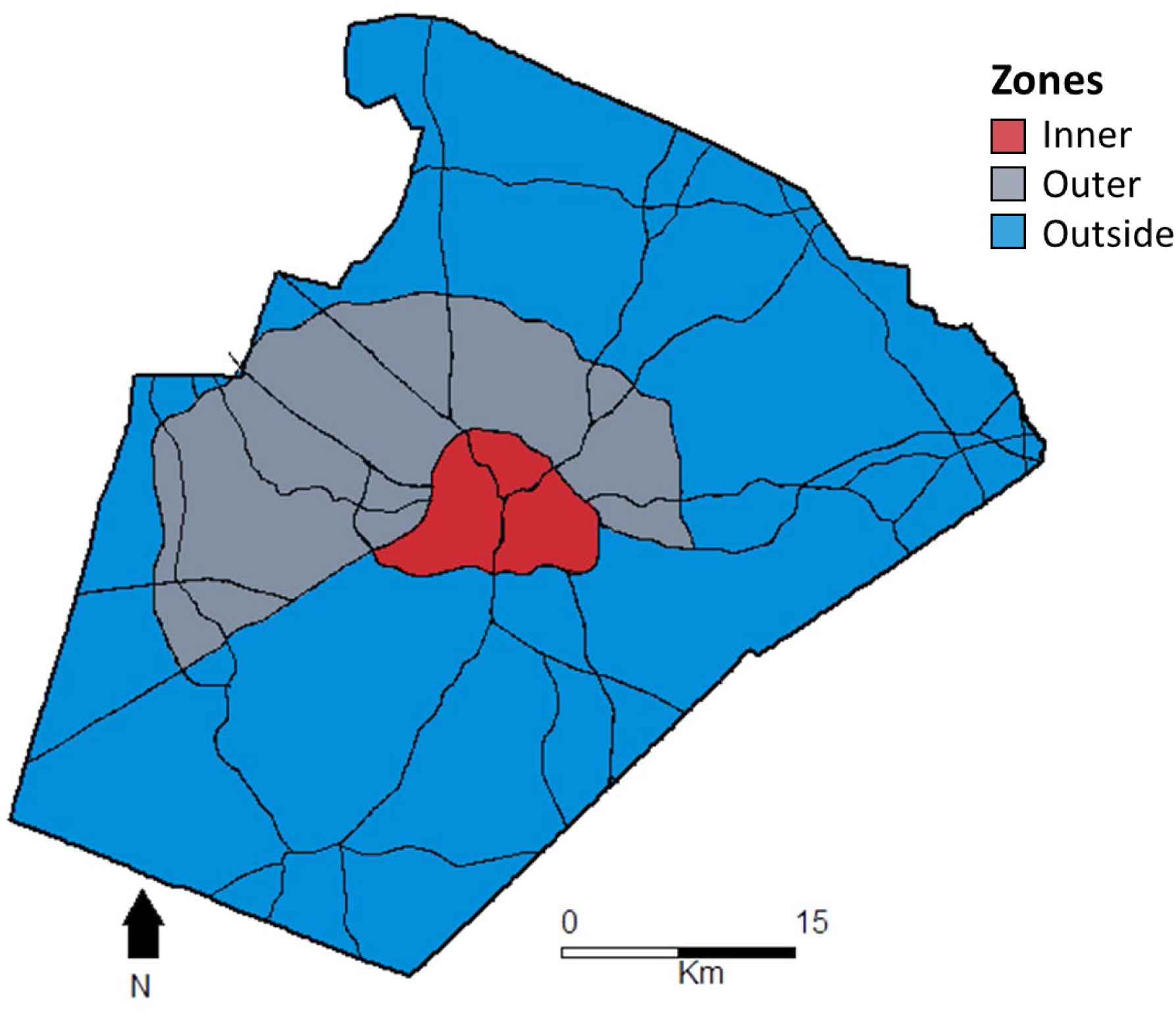
Delineation of zones for Wake County which act as proxies for the overall pattern of urbanization. In order from most to least urbanized: the Inner zone is defined as inside the interstate 440 beltline, the Outer zone is between the Interstate 540 beltline and I-440, and the Outside zone the remainder of the county.

### Sampling

We used *Ae. albopictus* samples collected from Wake County, North Carolina (Fig. 2). We sampled over two different years, 2016 and 2018. We collected *Ae. albopictus* eggs from fifteen sites in 2016 weekly between 15 April – 26 October 2016 as part of a state-wide mosquito survey effort to characterize the distribution and abundance of *Aedes* species in North Carolina (Reed et al. 2019; Fig. 2A). We placed traps strategically around the county in areas with some accumulation of human refuse and along forest/open edges. The fifteen locations were chosen for convenience and *Aedes* habitat suitability: five waste and recycling management centers, four gas stations, two residential backyards, and four miscellaneous buildings (school, government building, museum, retail). We collected eggs at each site using three ovitraps and hatched, reared, and identified the mosquitoes in the lab (for full methods, see Reed et al. 2019).

**Figure 2.**
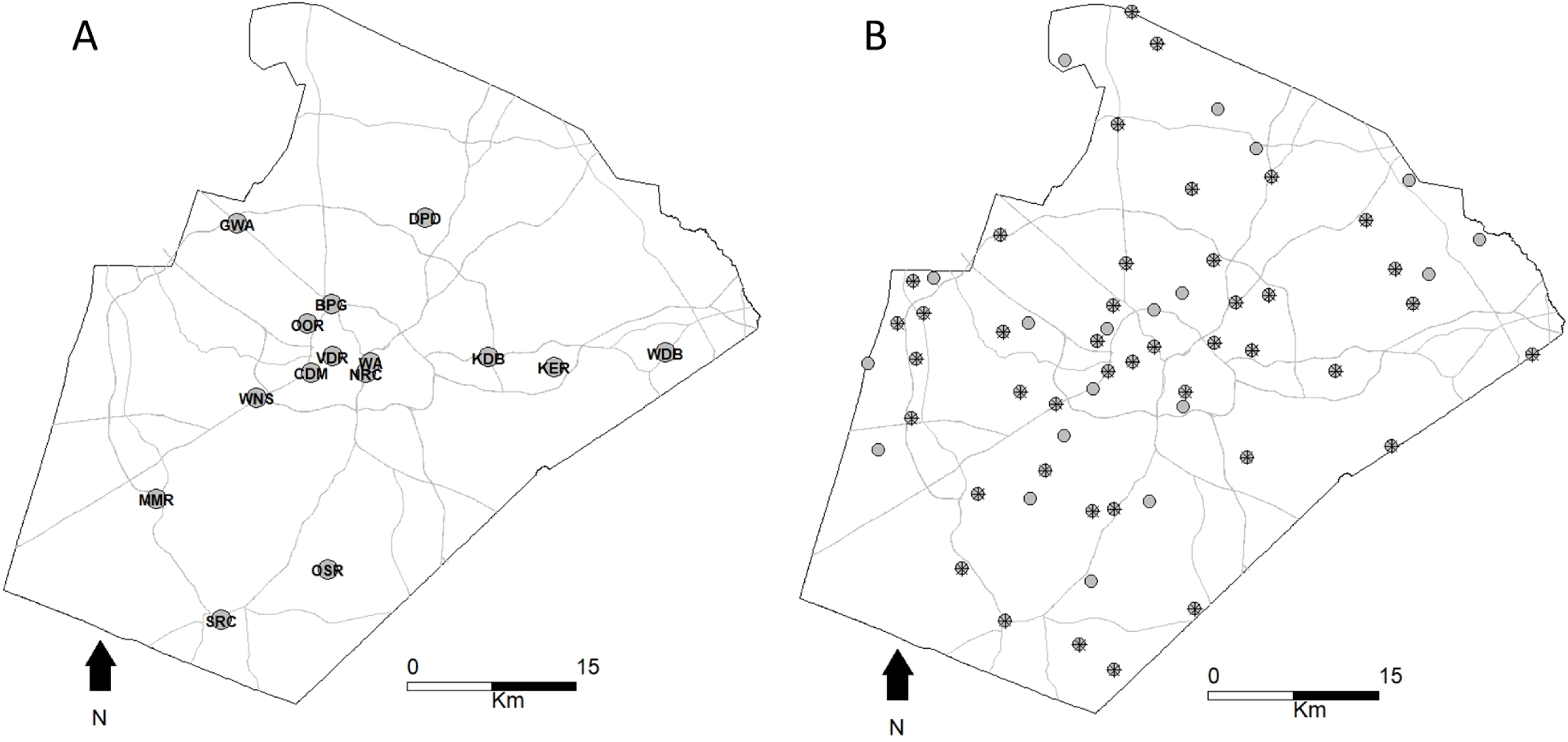
Wake County sampling locations. (A) sites and site names from 2016; all have genetic data for a portion of collected individuals. There are four sites in the Inner zone, five populations in the Outer zone, and six populations in the Outside zone. (B) Sampled sites from 2018; points overlaid with asterisks are sites that also have genetic data for mosquitoes. The number of sites in each zone (and number of sites used in population genetic analyses in parentheses) are as follows. Inner zone: six (four); Outer zone: 18 (14); Outside: 37 (24).

In 2018, we collected adult mosquitoes at 61 sites (Fig. 2B) using BG sentinels baited with BG Lures, a proprietary chemical attractant targeted at anthropophilic *Aedes* species (Biogents GmbH, Regensburg, Germany). Three of our 61 sites were identical to the sites in 2016. To determine the locations of the remaining sites, we used the *r.random.cells* function in GRASS GIS (GRASS Development Team, 2018) to generate 100 random points across Wake County, each with a 1000m buffer within which no other points could fall. We chose the 1000m buffer size because previous studies found that *Ae. albopictus* rarely disperse further than this distance (Honório et al. 2003, Medeiros et al. 2017). For each of these 100 points, we generated a 100 m buffer within which we could place a trap. We then selected points that were deemed “accessible,” which we defined as areas within 1 Km of a public road and that were not entirely water, which eliminated 21 potential sampling locations. Of the 79 remaining, we randomly selected 60 using a random number generator in RStudio (RStudio Team 2018). We then exported these points and their buffers to Google Earth (Google Earth, Google 2008). Using Street View, we ruled out 6 of these sites because the buffers fell completely within private lands. The remaining 56 points had buffers that fell at least partially within public land. After gaining permission to place traps with Wake County Department of Health and Human Services, we collected mosquitoes from 7 June – 25 June 2018, sampling each location once a week, leaving the traps out to collect mosquitoes for 24 hours. We then conducted an extra day of sampling for locations where a trap had failed, either due to a defective battery or removal of the collection funnel and bag (9 occurrences/181 trap*nights).

Given that we used different collection methods for the different years, and because we also found differences in genetic patterns between sampling types (Reed et al. 2023), we limited our comparisons within years for both abundance estimates and measures of genetic structure. To address the potential that egg sampling may bias towards greater genetic differentiation than adult sampling, we only used one to three individuals reared from each ovitrap for a given week.

### Genomic Library Preparation

We selected a subset of collected mosquitoes from both years to build genomic libraries for sequencing following Burford Reiskind et al. (2016), which uses a double-digest restriction-enzyme associated DNA (ddRAD) approach. We included sites in the final analysis that had more than three individuals with at least 8 ng DNA/µL after DNA extraction following the manufacturer’s protocol (Qiagen DNAeasy Tissue Kits), with one amendment, in which we incubated mosquitoes for an initial 24h period in Proteinase K. We used two restriction enzymes, *SphI* and *MluCI*, to fragment their genomes. We annealed barcodes to each fragment, size-selected fragments between 350-475 bp long using the BluPippin Prep (Sage Science) at the NCSU Genomic Sciences Laboratory. We built and sequenced 11 genomic libraries, each with 48 individuals for a total of 15 sampling sites with 192 individuals in 2016, and 42 sampling sites with 336 individuals in 2018. We completed all sequencing at the NCSU Genomic Sciences Laboratory on the Illumina HiSeq 2500 and we conducted single-end sequencing of 100 bp fragments.

### Bioinformatic Processing

The Illumina platform de-multiplexed to indices differentiating the two libraries in each lane, producing one FASTQ file per library. For each library, we checked the phred score to ensure high quality of sequence reads using FASTQC (Babraham Bioinformatics; http://www.bioinformatics.babraham.ac.uk/projects/fastqc). We then ran the *process_radtags* script in STACKS version 2.00 (Catchen et al. 2011) to filter low-quality reads (phred score < 33), trim reads to 90 bp, and demultiplex barcodes to produce FASTQ files for each individual following Burford Reiskind et al. (2016). We conducted one STACKS *denovo* pipeline with all individuals (*n* = 528), which generated a unique catalog of SNPs. We used the following parameters: (1) minimum read depth (*-m*) of six, (2) maximum number of mismatches between loci for an individual (*-M*) of 3, and (3) maximum number of mismatches allowed between loci for the catalog (*-n*) of 2. We ran STACKS *Populations* pipeline to remove SNPs that did not meet the following parameters: (1) present in a minimum of two sites (*-p* = 2) and (2) were not present in 75% of individuals in a population containing that SNP (*-r* = 0.75). We ran the *Populations* pipeline with two groups: (1) Wake 2016 individuals and (2) Wake 2018 individuals.

Following the *populations* pipeline, we further filtered SNPs using PLINK v.1.19 (Purcell et al. 2007), removing variants with a minimum allele frequency (MAF) of less than 0.01 and a genotyping rate (GENO) of less than 0.5. We then removed individuals that had less than 25% of the remaining SNPs genotyped (MIND). Finally, we used the *hw.test* function in the R package *pegas* v 0.15 (Paradis 2010) to identify SNPs out of Hardy-Weinberg Equilibrium and removed variants with a *P* value below the threshold value after applied a Bonferroni correction (*P* < 0.05 / # SNPs).

### Abundance Analyses

We tested significant differences in *Ae*. *albopictus* abundance between zones using a generalized linear mixed model assuming a negative binomial distribution covariance structure to account for natural aggregation in arthropod distributions in JMP 18.0 (2024, JMP Statistical Discovery LLC, Cary, NC). For 2016 data, we compared eggs per ovitrap*week in each of our 15 sites categorized into our three zones (inner, outer, outside). For 2018 data, we compared adult *Ae. albopictus* per daylight trap hour to correct for trapping effort in 62 sites and to reflect the diurnal activity of *Ae. albopictus* (Unlu et al 2021, Wynne et al. 2024), categorized into our three zones (inner, outer, outside).

### Genomic Analyses

We ran population genomic analyses to investigate patterns in genetic diversity, differentiation, and structure between sampling sites. To measure genetic diversity, we calculated expected heterozygosity (*H_E_*) and the inbreeding coefficient (*F_IS_*) corrected for small sample size for each location using the *genetic_diversity* function in the R package *gstudio* v1.5.2 (Dyer 2016). We evaluated significant differences in mean genetic diversity between zones using GLS, described above. To test for genetic differentiation, we estimated pairwise *F_ST_* among sampling sites for each year using the R package *hierfstat* v0.5-11 (*wcfst*; Goudet 2015) and tested significant differences in pairwise *F_ST_* using 1000 bootstraps and generated 95% confidence intervals (CIs; *boot.ppfst*). We determined significant differences when CIs did not include zero up to four significant figures.

We evaluated genetic structure in multiple ways. First, we measured isolation by distance among sample sites and across zones using a Mantel test (Sokal 1979). Second, we used the program STRUCTURE v.2.3.4 (Pritchard et al. 2000, Hubisz et al. 2009), which implements an individual-based Bayesian iterative algorithm to assign individuals to user-defined *k* clusters. We ran STRUCTURE using the admixture ancestry model with 10,000 burn-ins, 10,000 MCMC replications, and a *k* ranging from 1 to 10 with 10 iterations per *k* for the following datasets: 2016 individuals and 2018 individuals. We used STRUCTUREharvester (Earl and vonHoldt 2011) to determine the value of *k* with the highest likelihood using the Evanno method (Evanno *et al*., 2005). Third, we evaluated structure by conducting a discriminant analysis of principal components (DAPC), implemented in the R package *adegenet* v2.1.11 (Jombart 2008). DAPC can reveal more complex spatial genetic structure than k-mean clustering algorithms such as STRUCTURE and does not make assumptions based on population genetic models (Plue et al. 2018). We chose the optimal number of principal components (PCs) based on the PC value with the lowest root mean squared error and the highest mean success rate after cross-validation with a training set size of 0.95, 1000 replicates, and a maximum number of PCs equal to a third of the individuals included in the dataset to prevent overfitting the data (Jombart et al. 2008, Plue et al. 2018).

We further investigated population genetic differentiation in DAPC within and between zones in Wake County. To investigate genetic structure within zones, we used cross-validation in DAPC to decide the optimal number of PCs to retain. To look at differences in degree of differentiation between zones, we fit zones separately with the same number of PCs and we compared correct assignment rates. We interpreted higher rates as indicating greater genetic differentiation between sites within a given zone. To account for different numbers of sites within zones in 2016 and 2018, we used a rarefaction method to generate a frequency distribution for zones with more sites. To do this, we determined the zone with the fewest sites and randomly selected an equal number of sites from the remaining zones. We then found the correct assignment rate for that subset of populations. We repeated this process over 1000 iterations to generate the frequency distribution for the zone and from which we calculated a mean and 95% confidence interval.

## Results

### Mosquito Abundance

Over 80% of the mosquitoes trapped were *Ae*. *albopictus* (Supp. Table S1A). In 2016, the sites in the Outer zone between beltlines I-440 and I-540 had the highest average abundance with 110.88 ± 31.48 (SEM) eggs per ovitrap per week, followed by the Outside zone (97.61 ± 22.70 (SEM) eggs per trap per week) with the lowest in the Inner zone (65.37 ± 31.62 (SEM) eggs per trap per week). In spite of these differences, they were not significant in abundance between zones (*F* = 1.919, *df* = 2 *P* = 0.1929; Fig. 3A).

**Figure 3.**
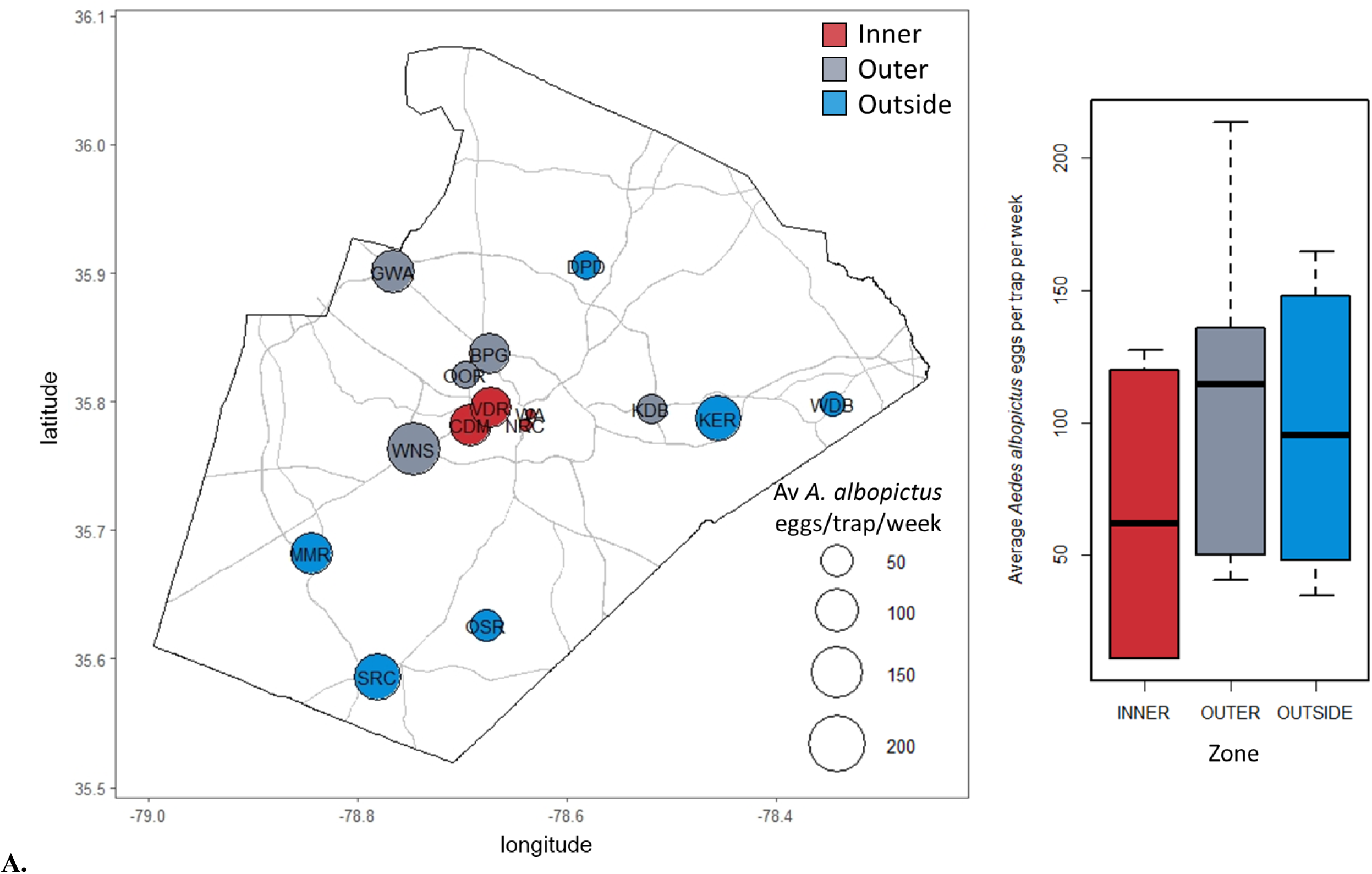
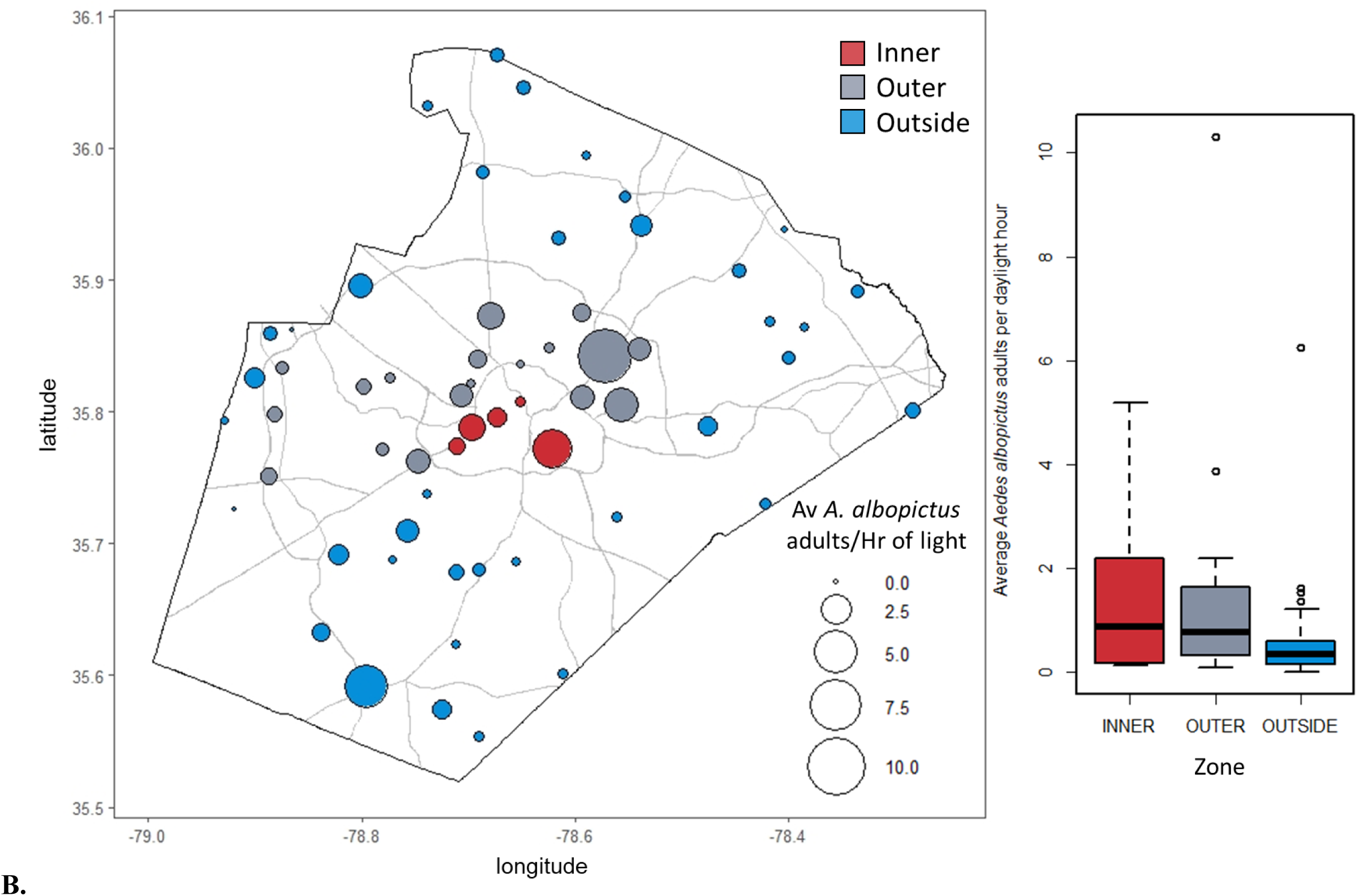
Average *A. albopictus* egg abundance per week in Wake County. (**A**) In 2016, larger points indicate higher average egg abundance. *Aedes albopictus* eggs were found at all sampling locations, and average count ranged from 10.75-213.59 per ovitrap per week. There were no significant differences in mean abundance between zones. Sites in the Inner zone are shown in red, those in the Outer zone in gray, and in the Outside zone, blue. (**B**) Average *A. albopictus* adult abundance per light-hour in Wake County 2018 at 61 locations. Larger points indicate higher average adult abundance. *Aedes albopictus* adults were found at 59/61 sites, and average abundance ranged from 0.02—10.31 adults per light-hour. There were no significant differences in mean abundance between zones. Sites in the Inner zone are shown in red, those in the Outer zone in gray, and in the Outside zone in blue.

In 2018, we trapped 2,553 mosquitoes, of which 2,086 were *Ae*. *albopictus* (81.71%; Supp. Table S1B). Of the 61 sites we sampled, we found *Ae*. *albopictus* at all but two (Fig. 3B). Overall, the trap rate per daylight hour, defined as the hours between sunrise and sunset on the day(s) of collection, of *Ae*. *albopictus* across all sites was 0.9757 ± 0.2143 adults/daylight hour. The highest average abundance was in the Inner zone, with 1.58 ± 0.79 (SEM) *Ae*. *albopictus* adults per hour of daylight, while the lowest average abundance was in the Outside zone with 0.612 ± 0.17 (SEM) *Ae*. *albopictus* adults per daylight hour across 44 sites. The Outer zone average was close to the Inner zone, with 1.53 ± 0.56 (SEM) adults trapped per daylight hour. As in 2016, there were no significant differences in adult abundance between zones (*F* = 2.113, *df* = 2, *P* = 0.1303).

### Population Genetics: 2016

In 2016, the STACKS *de novo* pipeline identified 4,003,920 SNPs across 192 individuals from 15 sites, and 166,852 were retained after the *populations* pipeline. During the filtering step SNPs were further culled for minimum allele frequency, genotyping rate, and HWE (Table 1). We removed eleven individuals for low genotyping rates, leaving a final dataset of 181 individuals and 28,347 SNPs after filtering (Table 1).

**Table 1.** Summary of sequencing and bioinformatic processing and single nucleotide polymorphism (SNP) filtering for two years of samples for Wake County, North Carolina: 2016 and 2018. Includes, the total number of sites, the range of individuals sequenced per site (*n*), total number of individuals and variants, and number of individuals and variants per each step of filtering from the *de novo* pipeline to the HWE filtering. N.B. In 2016, we limited sampling to 1-3 individuals for a given ovitrap*week to avoid sampling siblings.

| **Year** | **No. Sites** | **Range *n* per site** |  | **De novo Pipeline (STACKS)** | **Populations Pipeline (STACKS)** | **MAF (Plink)** | **GENO (Plink)** | **MIND (Plink)** | **HWE - Final Dataset** |
| --- | --- | --- | --- | --- | --- | --- | --- | --- | --- |
| **2016** | **15** | **7-18** | **Individuals** | 192 | 192 | 192 | 192 | 181 | 181 |
|  |  |  | **Variants** | 4003920 | 166853 | 151184 | 34695 | 34695 | 28347 |
| **2018** | **42** | **5-10** | **Individuals** | 336 | 289 | 289 | 289 | 276 | 276 |
|  |  |  | **Variants** | 3096027 | 183753 | 169713 | 16066 | 16066 | 11305 |

Expected heterozygosity of sites ranged from 0.1203 to0.1389 and mean *H_E_* did not differ between the Inner, Outer, and Outside zones (*P* = 0.4360). The inbreeding coefficient *F_IS_* ranged from −0.0250 to0.0953. Only one site, WA in the Inner Zone, had a negative *F_IS_* value, which indicates that individuals sampled at this location had higher levels of heterozygosity than expected under Hardy Weinberg Equilibrium. Zone did not have a significant effect on average *H_E_*, but zone had a significant effect on *F_IS_* (*P* = 0.0463), with the Outside Zone having a significantly higher mean *F_IS_* value (0.0826) than the Inner zone (0.0401; Table 3).

**Table 2.** Mean expected heterozygosity (*H_E_*) and inbreeding coefficient (*F_IS_*) within groups for each year pair in Wake County, Raleigh, NC. Asterix (*) indicates groups whose differences in genetic diversity were statistically significant.

| **Year** | **Zone** | **$H_E$** | **$F_{IS}$** |
| --- | --- | --- | --- |
| 2016 | Inner | 0.1300 | 0.0401* |
|  | Outer | 0.1350 | 0.0717 |
|  | Outside | 0.1320 | 0.0826* |
| 2018 | Inner | 0.0984 | 0.1120 |
|  | Outer | 0.1020 | 0.0790 |
|  | Outside | 0.1000 | 0.0850 |

**Table 3.**
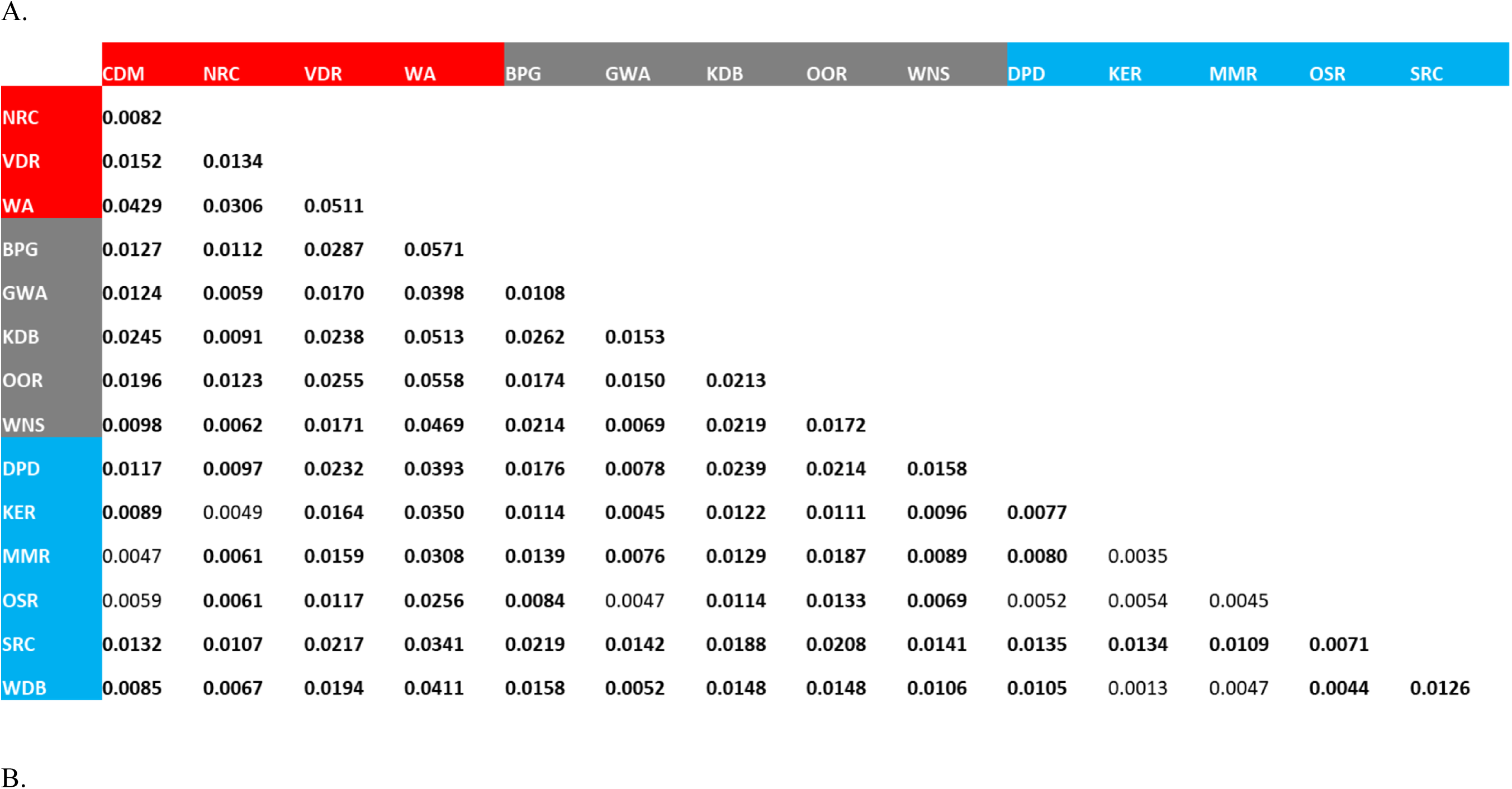
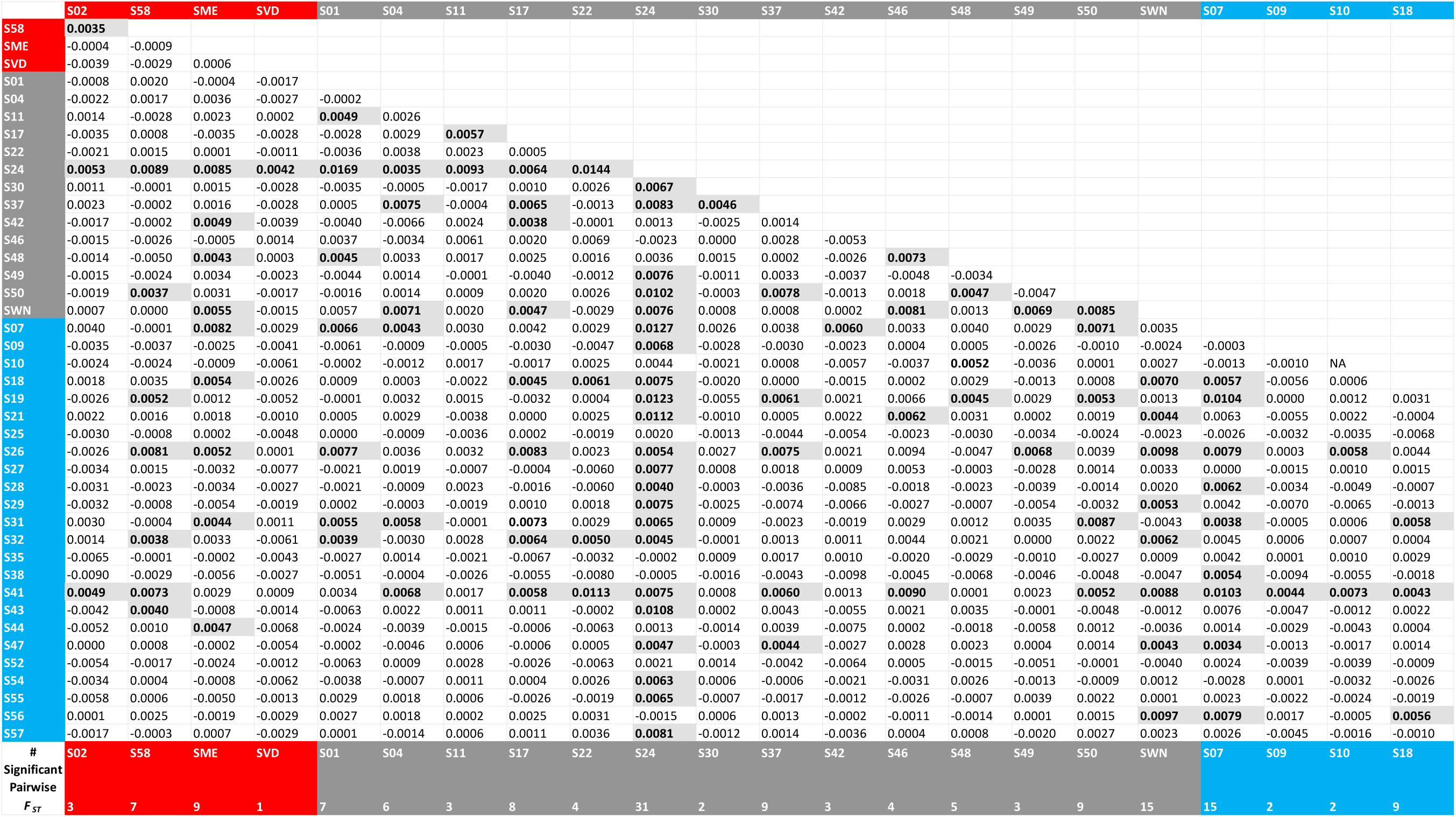
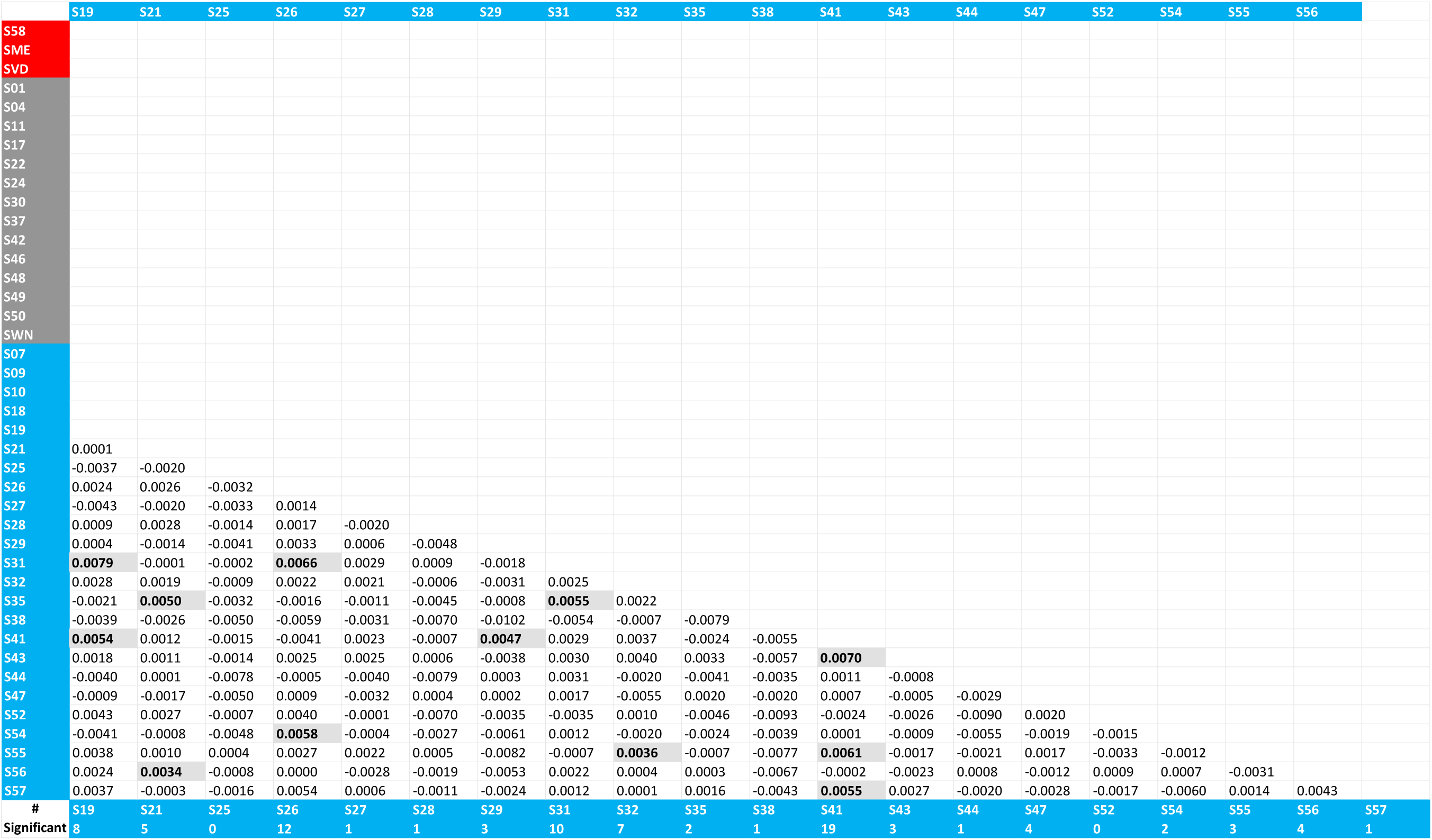
*Aedes albopictus* pairwise *F_ST_* values among sites in (A) Wake 2016 and, (B) Wake 2016 in the R package *hierfstat*. Background color of site names shows the zone that site belongs to: red = Inner, gray = Outer, and blue = Outside. Comparisons that were identified as significantly differentiated following a bootstrap analysis in *hierfstat* (bootstraps = 1000) generating 95% CIs. We restricted our significance test using CIs that did not include zero to three significant figures. Bolded numbers indicate significant pairwise comparisons using *hierfstat* bootstrap CIs. For the 2018 data set, the last row indicates the number of significant pairwise comparisons for each site. Two sites are identical between 2016 and 2018, WNS - SWN and VDR - SVD.

We found evidence of genetic differentiation in 2016 for many of the comparisons (*hierfstat F_ST_* values ranged from 0.0013 to 0.0571; Table 3) and these were significantly different from zero for 95 of the 105 comparisons. Two sites, VDR and WA, showed greater genetic differentiation compared to other locations than other sites in the data set. Both these sites were found in the Inner zone. WA was a restaurant located in a mixed residential/commercial area and VDR was a residential home in downtown Raleigh close to downtown Raleigh. To a lesser degree, the pairwise *F_ST_* results showed greater genetic differentiation at sites OOR, WNS, and KDB in the Outer zone. The site OOR was in a residential location just Outside the 440 beltline and the other two sites were located at Gas Stations in Cary and Knightdale respectively. The KDB site was adjacent to a large residential area, while the WNS site was adjacent to a shopping mall. Our analysis of overall genetic structure using STRUCTURE supported three clusters (*k*) using STRUCTUREharvester and the Evanno method (*k* = 3; Supp. Fig. S1A).

We retained 50 PCs after cross-validation with a correct assignment rate of 0.807 in DAPC. Sample sites primarily formed one cluster, with three sites, WNS and KBD (Outer Zone) and WA (Inner Zone), isolated from the remaining individuals (Fig. 4A). There was no signal of isolation by distance (Mantel test: observation = −0.3934; *P* = 0.977) across all sites. However, we did find evidence of isolation by distance in the Outside zone (Mantel test: observation = 0.3833; *P* = 0.01). For DAPC comparisons within zones, the Inner zone had 13 PCs retained after cross validation, which yielded a correct assignment rate of 0.9048, the Outer zone had eight PCs retained and a correct assignment rate of 0.6964, and the Outside zone had 26 PCs retained with a correct assignment rate of 0.7831 (Wake County 2016 Fig. 5). To detect if there were any significant differences in correct assignment rates between zones, we used 13 PCs for all zones. We found a correct assignment rate mean of 0.9048, 0.8380 (95% confidence interval: 0.7556 – 0.8889), and 0.6654 (95% confidence interval: 0.5472 – 0.7924) in the Inner, Outer, and Outside zones respectively. Within each zone there was less clustering of sites in the Inner zone than the other two zones. However, there were sites that did not group within the other two zones such as KDB in the Outer zone and DPD and SRC in the Outside zone.

**Figure 4.**
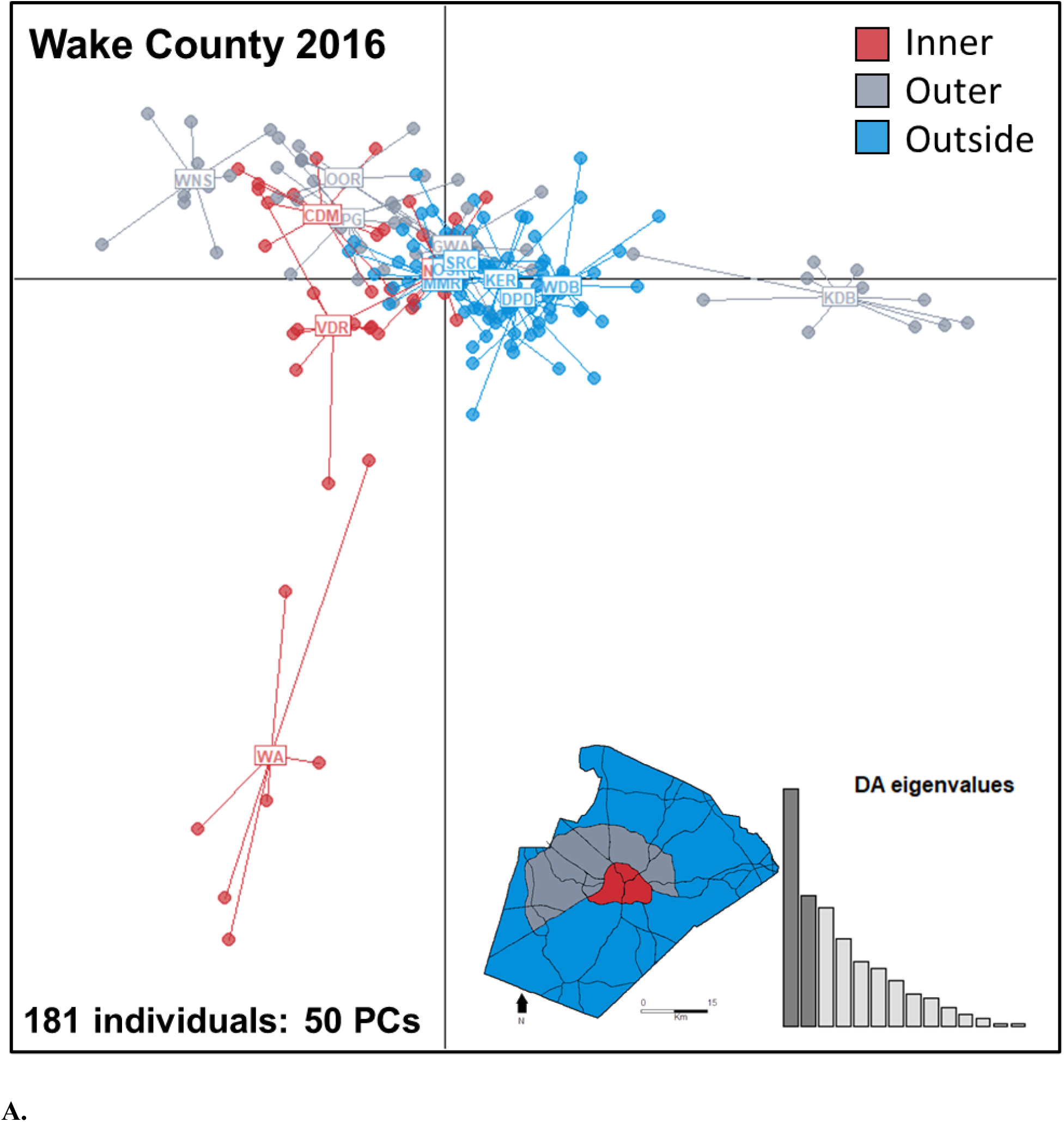
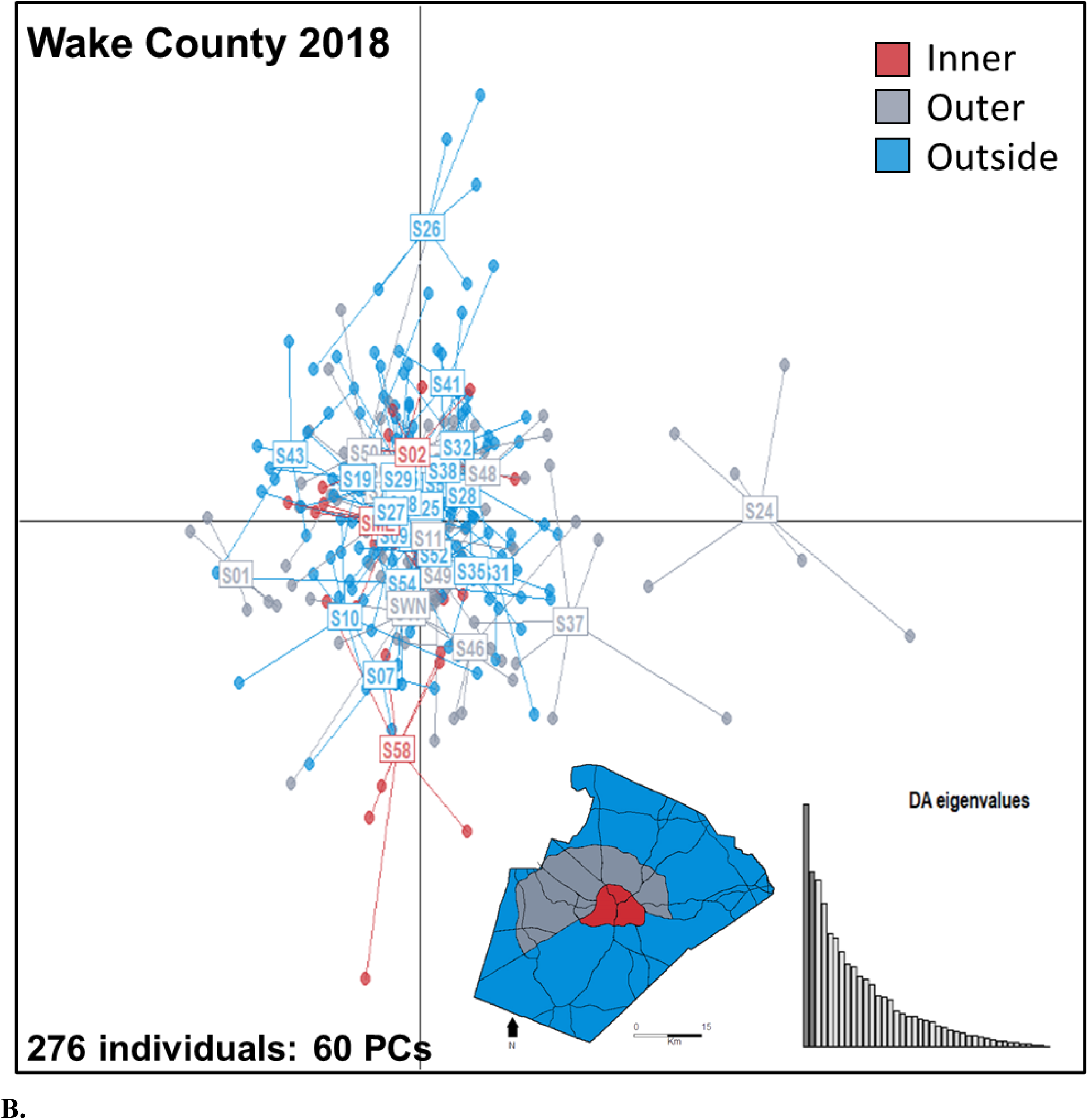
DAPC scatterplots for all sample sites. **(A)** 15 sites sampled in **Wake County 2016**, with the first two discriminant functions (DAs) on the *x* and *y* axes. Cross-validation retained 50 principal components, which produced a correct assignment rate of 0.8066. **(B)** for the 42 sites sampled in **Wake County 2018**, with the first two discriminant functions (DAs) on the *x* and *y* axes. Cross-validation retained 60 principal components, which produced a correct assignment rate of 0.6200

**Figure 5.**
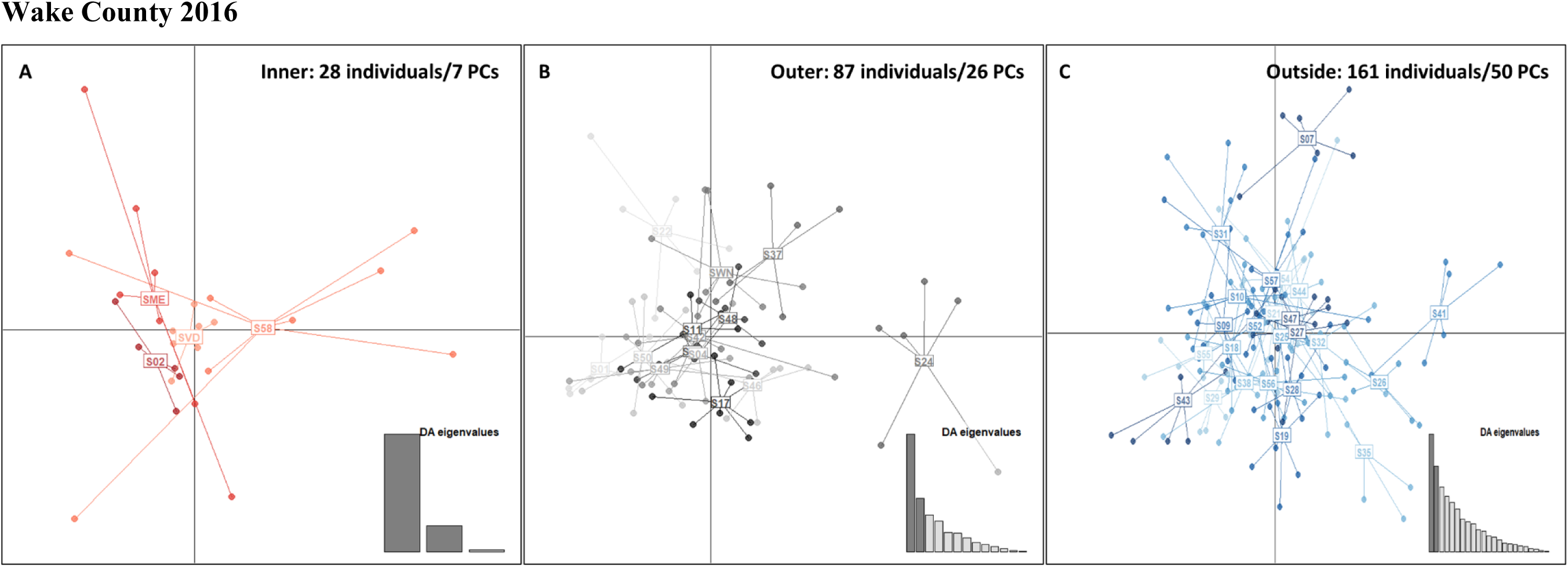
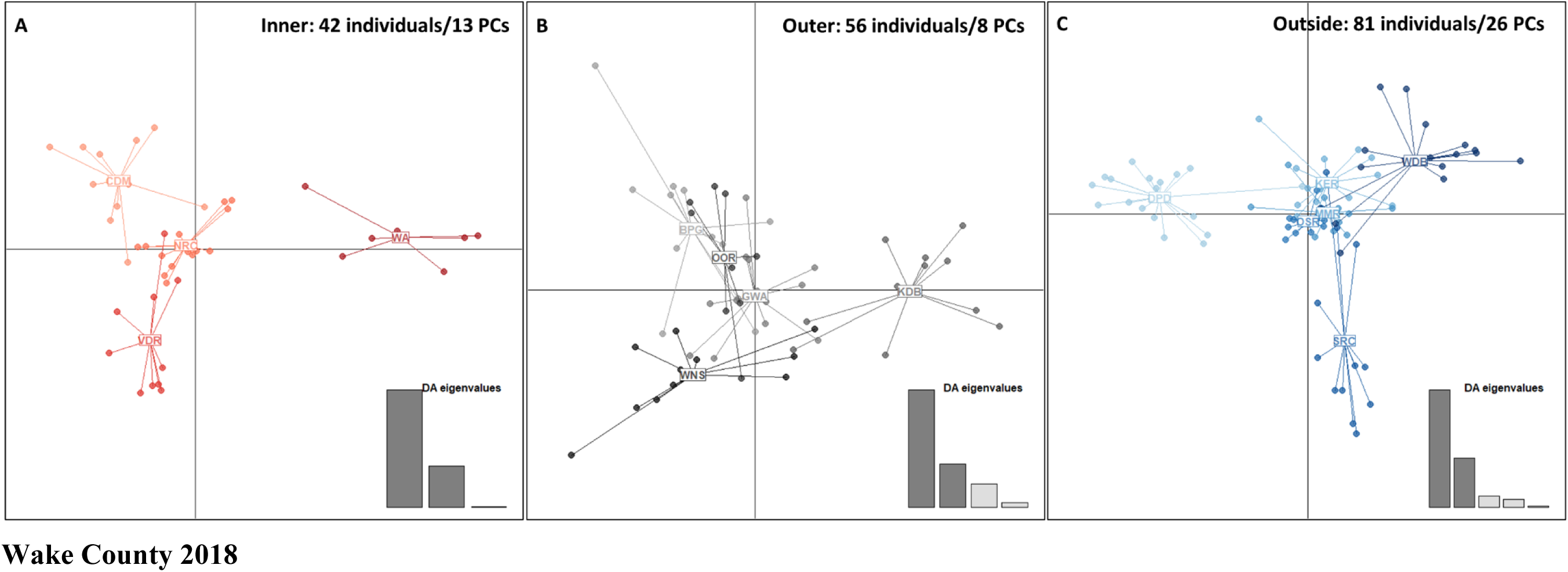
DAPC scatterplots for each zone of Wake County. **Wake county 2016**. The Inner zone had four populations with 42 individuals, the Outer zone had five populations with 56 individuals, and the Outside zone had six populations with 81 individuals. The *x* and *y* axes show the first two discriminant functions. After cross validation, the Inner zone (A) retained 13 principal components (PCs) with a correct assignment rate (CAR) of 0.9048, the Outer zone (B) retained eight PCs (CAR = 0.6964), and the Outside zone (C) retained 26 PCs (CAR = 0.7831). **Wake County 2018.** The Inner zone had four populations with 28 individuals, the Outer zone had 14 populations with 87 individuals, and the Outside zone had 24 populations with 161 individuals. The *x* and *y* axes show the first two discriminant functions. After cross validation, the Inner zone (A) retained seven principal components (PCs) with a correct assignment rate (CAR) of 0.6787, the Outer zone (B) retained 26 PCs (CAR = 0.7011), and the Outside zone (C) retained 50 PCs (CAR = 0.7826).

### Population Genetics: 2018

In 2018, the STACKS *de novo* pipeline identified 3,096,027 SNPs. 183,753 SNPs were retained in the Populations pipeline. During the filtering step SNPs were further culled for minimum allele frequency, genotyping rate, and HWE (Table 1), leaving a final dataset of 276 individuals and 11,305 SNPs (Table 1).

There was minimal variation in expected heterozygosity, which ranged from 0.0937 to 0.1058, and we did not find significant differences in mean *H_E_* between zones (*P* = 0.1019) (Table 2). *F_IS_* was more variable and ranged between 0.0201 to 0.1562. The Inner zone had the highest inbreeding coefficient (0.1120), but we did not find significant differences between zones (*P* = 0.3040; Table 2). We also found significant genetic differentiation among sampling sites in the pairwise *F_ST_* analysis, but no evidence of isolation by distance. Pairwise *F_ST_* values ranged from −0.0102 to 0.0169. Of the 861 pairwise comparisons, 425 (49.4%) had zero or negative *F_ST_* values to four significant figures. The five sites with the greatest number of significant pairwise *F_ST_* were S24, SWN, SO7, S26 and S41 with between 12 and 31 significant differences (Table 3B). Both S24 and SWN were in the Outer zone, with the remaining sites in the Outside zone. In general, all five sites showed significant pairwise differences that were not specific to a particular zone. We found greater genetic divergence among sites within the Outer and Outside zones than in the Inner zone, though there was a marked difference in the number of overall comparisons per zone. STRUCTUREharvester identified an optimal *k* =4, though most individuals had much of their ancestry assigned to the same cluster, which is a pattern consistent with a true value of *k* = 1 (Supp. Fig. S1B).

In DAPC, we retained 40 PCs after cross validation for all individuals, with a correct assignment rate of 0.6200 across 42 populations (Fig. 4B). Most populations formed a single cluster, with some differentiation from the major cluster at sites S24, S26, and to a lesser degree at sites S41, S58, S27 and S01. The two sites that were most differentiated, S24 and S26, were located in a wooded area between businesses and a residential community in the Outer zone and in an herbaceous area adjacent to croplands and residential single-family homes in the Outside zone, respectively. In comparison to the pairwise *F_ST_*, these results suggest six of the 42 populations were genetically differentiated (Fig. 4B).

When divided among Inner, Outer, and Outside zones (Fig. 5 Wake County 2018), after cross-validation, we retained seven PCs in the Inner zone with a correct assignment rate of 0.6786, 26 PCs in the Outer zone (correct assignment rate = 0.7011), and 50 PCs in the Outside zone (correct assignment rate = 0.7826). To compare correct assignment rates, we used rarefaction methods with the Outer and Outside zones and used seven PCs for all zones. The mean correct assignment rate for the Outer zone was 0.6008 (95% CI: 0.4583 – 0.7917) and 0.5169 for the Outside zone (95% CI: 0.3571 – 0.6786) compared to the baseline 0.6786 for the Inner zone. For the DAPC analysis within zones, the same locations that were Outside of the main cluster in Figure 4 were as well within zones, with the addition of S35 in the Outside zone (Fig. 6 Wake County 2018).

## Discussion

We found spatial genetic variation that was not consistent year to year, with greater genetic divergences in 2016 than in 2018. Broadly, we found that mean egg abundance and expected heterozygosity was lowest in sites within urban zones, indicating a loss of genetic diversity, but populations sampled in rural zones were more inbred. We also observed more genetic structure among sites within urban zones than transitioning or rural zones in Wake County, in 2016. However, this pattern was not as clear in 2018 where we found genetic structure in all three zones biased towards the Outer and Outside zones where we had more sites. The results from the DAPC analysis also supported spatial variation in both years and mirrored the patterns we saw in the pairwise *F_ST_* results. However, the pattern of greater genetic differentiation within the Inner zone in 2016 was not identical in 2018. The dynamics here suggest that there is variation year to year and among sites that may be influenced by abiotic factors and small population dynamics at the scale of an urban to rural landscape.

Trends in abundance in Wake County within each year between 2016 eggs and 2018 adults were somewhat contradictory. Specifically, the Inner zone had the lowest egg abundance relative to other sites in 2016 and the highest adult abundance in 2018, though differences were not statistically significant in either year. While the scope of this study does not permit us to identify whether this switch was due to changes in mosquito population dynamics, land-use changes, or sampling methods between years, sampling methods may be the most logical explanation. Sampling method could also drive the greater genetic differentiation among sites in 2016 compared to 2018, as was highlighted in a within year study that compared egg and adult samples from the same locations in Wake County (Reed et al. 2023). Targeting the egg-stage is most effective when there are fewer competing larval habitats, while targeting host seeking adults is most effective when there are fewer hosts (Silver, 1989). These two situations may differ by level of urbanization in different ways, affecting ovitrapping and adult trapping differently. Our site selection might also influence this. In 2016, sites were specifically chosen based on convenience and where mosquitoes were likely to be found, while in 2018 sample sites were chosen to capture a wider area, with most sites chosen randomly. Many of the sites in the 2018 Outside zone were in landscapes less hospitable to high densities of *Ae*. *albopictus*. For example, some sites were located by agricultural fields, in the forest interior, and in high grass, all areas not associated with high *Ae*. *albopictus* abundance (Barker et al. 2003, Reiskind et al. 2017). These results are also consistent with rural land use bifurcation (Warren et al. 2018). This may explain why we did not see the same urban/suburban difference in abundance in 2018.

The population genetic analyses suggest variable genetic structuring between years and in strength. The expected heterozygosity was lowest at sites in the most urbanized zone (Inner zone) and highest in the intermediate zone (Outer zone). However, these differences were not statistically significant. These results suggest genetic diversity may be lower for *Ae*. *albopictus* in built landscapes and could indicate a recent population bottleneck and genetic isolation. This is similar to Manica et al.’s (2016) finding that highly developed areas were not high-quality habitats for *Ae*. *albopictus*.

In contrast, patterns of mean inbreeding coefficients (*F_IS_*) were more variable. The inbreeding coefficient, despite its name, is not a direct measure of relatedness between individuals. Instead, it indicates the degree to which observed levels of heterozygosity within a population deviate from the expected levels of heterozygosity given the population’s allele frequencies at the genetic markers under study. If we consider two populations with the same level of expected heterozygosity, the population with the higher *F_IS_* value will have more inter-individual genetic variation, and the population with lower *F_IS_* will have more variation within the individual. Therefore, the relatively low levels of expected heterozygosity paired with low *F_IS_* in urbanized zones indicate that there were fewer biallelic SNPs in those populations, but that individuals were more likely to be heterozygous at those loci. This could indicate frequent mating events between unrelated individuals, perhaps from migration events or admixture of multiple bottlenecks. Overall, the patterns in the inbreeding coefficient suggest local, small-scale dynamics that drive population dynamics within each zone.

Between years, there was little consistency in genetic differentiation. For instance, we observed significant genetic structure across the county in 2016 in both the pairwise *F_ST_* and DAPC analyses, but lower genetic structure among sites in both analyses in 2018. However, the significant genetic structure in 2016 and 2018 were not reflected in the STRUCTURE results, which did not show strong differentiation among sites in either year. Based on the DAPC analysis, the Inner zone was most genetically differentiated, followed by the Outer and Outside zones in 2016, indicating little gene flow between the urban-most locations. For instance, in 2016, WA formed a separate genetic cluster in both DAPC scatterplots, despite being located less than 1000m from another sampled site, NRC. However, this pattern was not as strong in 2018 as it was in 2016, with all three zones showing similar degree of genetic differentiation. This suggests that in some years there is minimal gene flow between more urban locations but does not preclude in both years gene flow in or out of rural areas to more urban populations.

In 2018, we found five sites, two in the Outer and three in the Outside zones, that were differentiated from most other sites. These same sites also did not cluster within their zones in the DAPC analysis. While this was not at the same extent as we saw in 2016, it does suggest there is spatial heterogeneity across Wake County and potentially sources and sinks that vary year to year. We sampled a larger geographic area of Wake County in 2018 and perhaps have a more accurate estimate of the degree of differentiation among sites by using adults (Reed et al. 2023). However, in this geographic area there may be differences in degree of connectivity year to year, with some years supporting more widespread gene flow and lower genetic differentiation driven by factors we did not measure. While it was outside the scope of this study, a landscape genetic analysis would aid in better understanding source and sink dynamics and gene flow among these three zones.

The patterns we observed point to several areas for future research. First, we saw more genetic differentiation and variation in abundance when we sampled eggs rather than adults, even when we limited the number of individuals from a given ovi-trap*week. Many population and landscape genetic studies on *Ae*. *albopictus* use egg or larval samples, and these should be interpreted with caution (e.g., Ayres et al. 2002, Schmidt et al. 2017, Adilah-Amrannudin et al. 2018, Reed et al. 2023). If these sampling methods are used in research designed to inform mosquito control, especially those targeted for adult mosquitoes, overestimating genetic differentiation may lead to the misidentification of population management units.

Finally, this study emphasizes the challenging nuances of urban ecology, especially for nonnative species. We found evidence both consistent with and in opposition to the traditional urban—rural ecological paradigm (McDonnell and Pickett 1990, McDonnell et al. 1997). While the genomic approach we used is powerful, the spatio-temporal scale of our study presented challenges identifying genetic breaks in a region of potentially high gene flow and will require a finer landscape-genetic approach with more temporal, geographic, and genomic sampling. In addition, considering the increase in genetic control methods of mosquito control, additional studies should measure gene flow and directional migration between populations, especially between and within urban and rural locations, and investigate the role of the intervening landscape matrix in determining genetic diversity and connectivity. We did note genetic differentiation was generally highest in the most urbanized areas, but genetic diversity was overall similar across zones and patterns of abundance varied between years. Careful consideration of species ecology, and location-specific features of the urban—rural network, with intensive sampling, as well as a population genetic approach are necessary to predict population responses to human development and to inform management during disease outbreaks.

## Acknowledgements

We would like to thank Anastasia Figurskey, Allison Cousins and Chris Intehar for their assistance in field collection and identification, Emma Wallace for her help with DNA extractions and genomic library preparation, and Paul Labadie for consultation on molecular methods and bioinformatic processing. We also thank the Wake County Department of Health and Human Services for their permission to sample on public lands. This study was supported by the USGS Southeast Climate Adaptation Science Center graduate fellowship awarded to Emily M. X. Reed and was funded by the Wynne Innovation Grant from the CAL Dean’s Enrichment Grant Program at NCSU awarded to Martha O. Burford Reiskind.

## Author Contributions

Emily M. X. Reed: Conceptualization, Methodology, Data Collection, Formal analysis, and Writing-original draft. Michael H. Reiskind: Conceptualization, Methodology, Writing-reviewing and editing. Martha O. Burford Reiskind: Conceptualization, Methodology, Formal analysis, Writing-reviewing and editing.

