## Supplemental Table and Figure for "Spatiotemporal variation in abundance and genetic structure across the urban-rural landscape gradient: *Aedes albopictus* (Skuse, 1894) (Diptera: Culicidae) in Wake County, NC"

**Supplemental Tables**

**Table S1.** Abundance summary for all sampled *Aedes*-container mosquitoes in Wake County (**A**) 2016 and (**B**) 2018. Includes collection site, zone (Inside 1, Outer 2, Outside 3) total number (N) of egg papers or traps, eggs (N), total numbers of individuals (N), proportion of individuals (P), and estimated averages (A) for Albo (*Aedes albopictus*), Tris (*Ae. triseriatus*), and Japon (*Ae. japonicus*) for 2016. For 2018 includes collection site, site label, trap days, trap time, light time total (for total trap light time), total numbers N, proportion P for Albo (*Ae. albopictus*) and Other (*Aedes* other).

**A.**

| Site | Zone | Papers N | Eggs N | Albo N | Tris N | Japon N | Albo P | Tris P | Japon P | Albo A | Tris A | Japon A |
|---|---|---|---|---|---|---|---|---|---|---|---|---|
| BPG | 2 | 31 | 3553 | 1035 | 0.0000 | 0.0000 | 1.0000 | 0.0000 | 0.0000 | 114.6129 | 0.0000 | 0.0000 |
| CDM | 1 | 33 | 4229 | 1015 | 5.0000 | 0.0000 | 0.9951 | 0.0049 | 0.0000 | 127.5233 | 0.6282 | 0.0000 |
| DPD | 3 | 32 | 1539 | 421 | 0.0000 | 0.0000 | 1.0000 | 0.0000 | 0.0000 | 48.0938 | 0.0000 | 0.0000 |
| GWA | 2 | 33 | 4744 | 906 | 46.0000 | 9.0000 | 0.9428 | 0.0479 | 0.0094 | 135.5300 | 6.8812 | 1.3463 |
| KDB | 2 | 32 | 1626 | 678 | 7.0000 | 0.0000 | 0.9898 | 0.0102 | 0.0000 | 50.2932 | 0.5193 | 0.0000 |
| KER | 3 | 27 | 3989 | 676 | 0.0000 | 0.0000 | 1.0000 | 0.0000 | 0.0000 | 147.7407 | 0.0000 | 0.0000 |
| MMR | 3 | 31 | 4211 | 572 | 32.0000 | 6.0000 | 0.9377 | 0.0525 | 0.0098 | 127.3766 | 7.1260 | 1.3361 |
| NRC | 1 | 32 | 351 | 181 | 0.0000 | 0.0000 | 1.0000 | 0.0000 | 0.0000 | 10.9688 | 0.0000 | 0.0000 |
| OOR | 2 | 33 | 1337 | 479 | 2.0000 | 0.0000 | 0.9958 | 0.0042 | 0.0000 | 40.3467 | 0.1685 | 0.0000 |
| OSR | 3 | 33 | 2154 | 504 | 17.0000 | 0.0000 | 0.9674 | 0.0326 | 0.0000 | 63.1429 | 2.1298 | 0.0000 |

| | | | | | | | | | | | | |
|---|---|---|---|---|---|---|---|---|---|---|---|---|
| SRC | 3 | 32 | 5422 | 871 | 26.0000 | 0.0000 | 0.9710 | 0.0290 | 0.0000 | 164.5263 | 4.9112 | 0.0000 |
| VDR | 1 | 32 | 3591 | 920 | 0.0000 | 0.0000 | 1.0000 | 0.0000 | 0.0000 | 112.2188 | 0.0000 | 0.0000 |
| WA | 1 | 32 | 344 | 55 | 0.0000 | 0.0000 | 1.0000 | 0.0000 | 0.0000 | 10.7500 | 0.0000 | 0.0000 |
| WDB | 3 | 32 | 1165 | 192 | 9.0000 | 0.0000 | 0.9552 | 0.0448 | 0.0000 | 34.7761 | 1.6301 | 0.0000 |
| WNS | 2 | 32 | 6835 | 1284 | 0.0000 | 0.0000 | 1.0000 | 0.0000 | 0.0000 | 213.5938 | 0.0000 | 0.0000 |

**B.**

| **Site** | **Label** | **Zone** | **TrapDays** | **TrapTime** | **TrapLight** | **Albo N** | **Other N** | **Albo P** | **Other P** |
|---|---|---|---|---|---|---|---|---|---|
| S01 | BREN | 2 | 3 | 61.32 | 35.97 | 62 | 18 | 0.7750 | 0.2250 |
| S02 | STON | 1 | 3 | 64.98 | 40.48 | 7 | 1 | 0.8750 | 0.1250 |
| S03 | CARP | 3 | 3 | 65.7 | 40.35 | 4 | 1 | 0.8000 | 0.2000 |
| S04 | KIFA | 2 | 3 | 66.52 | 42.02 | 16 | 28 | 0.3636 | 0.6364 |
| S06 | CHFA | 3 | 3 | 63.12 | 37.85 | 3 | 7 | 0.3000 | 0.7000 |
| S07 | BOTH | 3 | 3 | 71.83 | 46.48 | 75 | 6 | 0.9259 | 0.0741 |
| S08 | OLCH | 3 | 3 | 53.05 | 27.7 | 0 | 2 | 0.0000 | 1.0000 |
| S09 | CREE | 3 | 4 | 90.93 | 57.15 | 21 | 18 | 0.5385 | 0.4615 |
| S10 | MPCR | 3 | 3 | 63.28 | 38 | 6 | 24 | 0.2000 | 0.8000 |
| S11 | MAVI | 2 | 3 | 56.47 | 31.1 | 25 | 4 | 0.8621 | 0.1379 |

| | | | | | | | | | |
|---|---|---|---|---|---|---|---|---|---|
| S12 | MORR | 3 | 3 | 62.53 | 37.22 | 2 | 9 | 0.1818 | 0.8182 |
| S14 | SSTA | 1 | 2 | 38.47 | 21.58 | 3 | 9 | 0.2500 | 0.7500 |
| S15 | REPA | 2 | 3 | 55.65 | 31.13 | 5 | 6 | 0.4545 | 0.5455 |
| S17 | ELLS | 2 | 3 | 59.32 | 34.05 | 132 | 4 | 0.9706 | 0.0294 |
| S18 | EYOU | 3 | 3 | 58.9 | 33.63 | 14 | 22 | 0.3889 | 0.6111 |
| S19 | PEAR | 3 | 3 | 68.23 | 42.88 | 8 | 0 | 1.0000 | 0.0000 |
| S20 | CBEB | 2 | 3 | 61.78 | 36.43 | 3 | 4 | 0.4286 | 0.5714 |
| S21 | PGWR | 3 | 3 | 57.03 | 31.67 | 26 | 5 | 0.8387 | 0.1613 |
| S22 | LOUI | 2 | 3 | 56.7 | 31.32 | 323 | 3 | 0.9908 | 0.0092 |
| S23 | PURD | 2 | 3 | 61 | 35.63 | 6 | 3 | 0.6667 | 0.3333 |
| S24 | NWCP | 2 | 3 | 65.9 | 40.68 | 24 | 15 | 0.6154 | 0.3846 |
| S25 | MIBE | 3 | 3 | 63.72 | 38.35 | 19 | 4 | 0.8261 | 0.1739 |
| S26 | BRPE | 3 | 3 | 62.53 | 37.18 | 7 | 3 | 0.7000 | 0.3000 |
| S27 | BROU | 3 | 3 | 55.92 | 30.53 | 6 | 9 | 0.4000 | 0.6000 |
| S28 | DOVE | 3 | 3 | 65.93 | 40.57 | 18 | 11 | 0.6207 | 0.3793 |
| S29 | STAR | 3 | 3 | 62.83 | 37.53 | 51 | 4 | 0.9273 | 0.0727 |
| S30 | LORI | 2 | 3 | 63.08 | 37.75 | 12 | 13 | 0.4800 | 0.5200 |
| S31 | TREG | 3 | 3 | 58.6 | 33.23 | 14 | 2 | 0.8750 | 0.1250 |
| S32 | SCPE | 3 | 3 | 58.5 | 33.1 | 36 | 5 | 0.8780 | 0.1220 |
| S33 | JAFA | 3 | 3 | 59.27 | 33.93 | 12 | 4 | 0.7500 | 0.2500 |

| | | | | | | | | | |
|---|---|---|---|---|---|---|---|---|---|
| S34 | OLST | 3 | 3 | 62.25 | 36.88 | 5 | 54 | 0.0847 | 0.9153 |
| S35 | KICR | 3 | 3 | 60.75 | 35.42 | 16 | 12 | 0.5714 | 0.4286 |
| S36 | WAPL | 3 | 3 | 62.7 | 37.32 | 8 | 3 | 0.7273 | 0.2727 |
| S37 | GLCR | 2 | 3 | 63.17 | 37.83 | 28 | 5 | 0.8485 | 0.1515 |
| S38 | HANK | 3 | 3 | 60.9 | 35.53 | 13 | 1 | 0.9286 | 0.0714 |
| S39 | HASE | 3 | 3 | 68.78 | 43.42 | 5 | 3 | 0.6250 | 0.3750 |
| S40 | OGC2 | 3 | 3 | 74.45 | 49.05 | 8 | 11 | 0.4211 | 0.5789 |
| S41 | NMAI | 3 | 2 | 44.98 | 28.13 | 176 | 16 | 0.9167 | 0.0833 |
| S42 | SEFA | 2 | 3 | 61.57 | 36.22 | 19 | 1 | 0.9500 | 0.0500 |
| S43 | KEME | 3 | 3 | 56.35 | 31.43 | 30 | 4 | 0.8824 | 0.1176 |
| S44 | QUCR | 3 | 3 | 71.72 | 46.37 | 44 | 8 | 0.8462 | 0.1538 |
| S45 | TREX | 3 | 3 | 64.37 | 38.97 | 0 | 5 | 0.0000 | 1.0000 |
| S46 | STST | 2 | 2 | 48.87 | 32.02 | 49 | 1 | 0.9800 | 0.0200 |
| S47 | FAYE | 3 | 3 | 60.97 | 35.62 | 11 | 0 | 1.0000 | 0.0000 |
| S48 | PIVI | 2 | 3 | 73.75 | 48.37 | 41 | 14 | 0.7455 | 0.2545 |
| S49 | LYNN | 2 | 2 | 43.88 | 27.05 | 59 | 2 | 0.9672 | 0.0328 |
| S50 | SURI | 2 | 3 | 63.93 | 38.58 | 58 | 1 | 0.9831 | 0.0169 |
| S51 | APWI | 3 | 3 | 60.55 | 35.2 | 5 | 0 | 1.0000 | 0.0000 |
| S52 | BRBR | 3 | 3 | 60.27 | 34.93 | 14 | 13 | 0.5185 | 0.4815 |
| S53 | BASI | 3 | 3 | 59.9 | 34.5 | 1 | 17 | 0.0556 | 0.9444 |

| | | | | | | | | | |
|---|---|---|---|---|---|---|---|---|---|
| S54 | FOXB | 3 | 3 | 66.7 | 41.32 | 25 | 5 | 0.8333 | 0.1667 |
| S55 | CHAS | 3 | 3 | 58.95 | 33.6 | 41 | 6 | 0.8723 | 0.1277 |
| S56 | PENN | 3 | 2 | 40.83 | 24.03 | 37 | 1 | 0.9737 | 0.0263 |
| S57 | BISS | 3 | 3 | 61.57 | 36.2 | 10 | 2 | 0.8333 | 0.1667 |
| S58 | ELEN | 1 | 3 | 59.08 | 33.7 | 175 | 14 | 0.9259 | 0.0741 |
| S60 | HOCR | 3 | 3 | 66.73 | 41.38 | 5 | 7 | 0.4167 | 0.5833 |
| SME | MET | 1 | 3 | 75.23 | 50.23 | 110 | 5 | 0.9565 | 0.0435 |
| SOO | OOR | 2 | 3 | 75.6 | 50.23 | 5 | 5 | 0.5000 | 0.5000 |
| SPV | PVD | 1 | 1 | 23.82 | 15.4 | 11 | 0 | 1.0000 | 0.0000 |
| SVD | VDR | 1 | 3 | 74.73 | 49.87 | 52 | 4 | 0.9286 | 0.0714 |
| SWN | WNS | 2 | 3 | 77.03 | 52.12 | 85 | 3 | 0.9659 | 0.0341 |

### Supplemental Figures

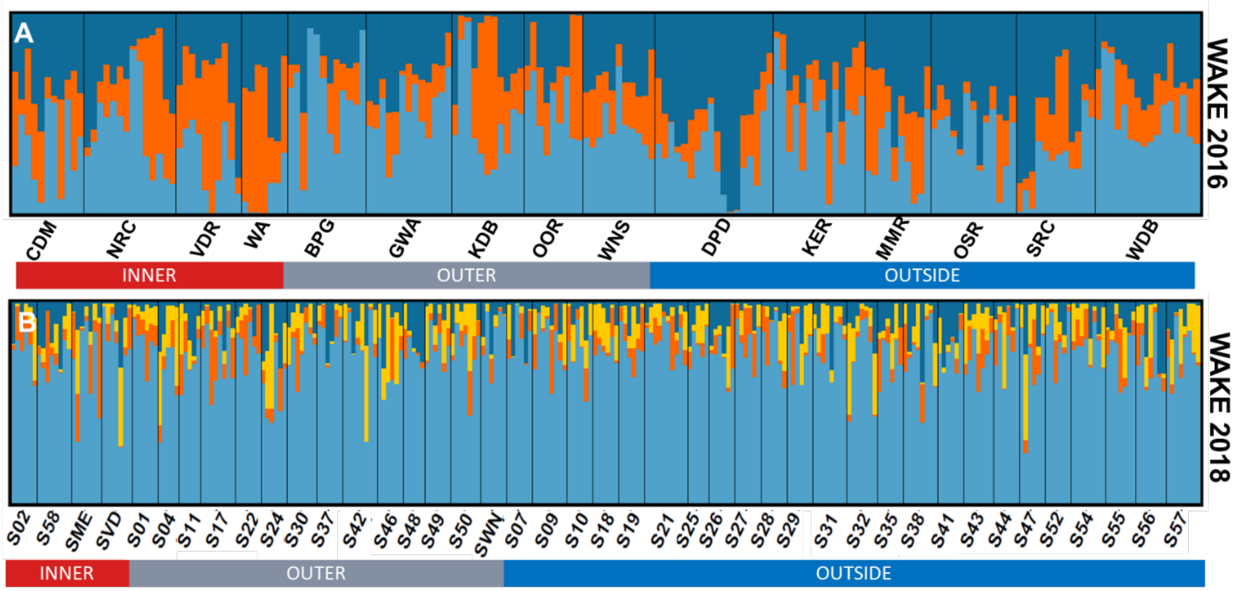


**Figure S1.** STRUCTURE bar plots with the optimal value of *k* determined by the Evanno method in STRUCTUREharvester for (A) Wake County 2016 ($k = 3$); and (B) Wake County 2018 ($k = 4$). Bar plots were generated using DISTRUCT v.
